# BCI learning phenomena can be explained by gradient-based optimization

**DOI:** 10.1101/2022.12.08.519453

**Authors:** Peter C. Humphreys, Kayvon Daie, Karel Svoboda, Matthew Botvinick, Timothy P. Lillicrap

## Abstract

Brain-computer interface (BCI) experiments have shown that animals are able to adapt their recorded neural activity in order to receive reward. Recent studies have highlighted two phenomena. First, the speed at which a BCI task can be learned is dependent on how closely the required neural activity aligns with pre-existing activity patterns: learning “out-of-manifold” tasks is slower than “in-manifold” tasks. Second, learning happens by “re-association”: the overall distribution of neural activity patterns does not change significantly during task learning. These phenomena have been presented as distinctive aspects of BCI learning. Here we show, using simulations and theoretical analysis, that both phenomena result from the simple assumption that behaviour and representations are improved via gradient-based algorithms. We invoke Occam’s Razor to suggest that this straightforward explanation should be preferred when accounting for these experimental observations.

## Introduction

Mammalian brains are remarkably robust and effective learning machines. They are capable of solving a bewildering variety of tasks while flexibly adapting to changing circumstances. Identifying the neural algorithms that support this learning is a critical step in developing our understanding of the brain (Richards et al., 2019; Rumelhart et al., 1988). However, it is challenging to study learning dynamics in typical behavioural tasks due to the complex relationship between a neuron’s activity and animal behaviour, which is ultimately what is rewarded.

In brain-computer interface (BCI) experiments (Fig 1a, Fetz (1969); Golub et al. (2018); Koralek et al. (2012, 2013); Oby et al. (2019); Sadtler et al. (2014); Santhanam et al. (2006)), neural activity itself is used to define an interface through which an animal must solve a behavioural task. This establishes a more direct and controllable pathway between activity and reward, providing a compelling setting for the investigation of neural learning dynamics and credit assignment (Athalye et al., 2020; Golub et al., 2016; Orsborn and Pesaran, 2017; Portes et al., 2022).

**Figure 1.**
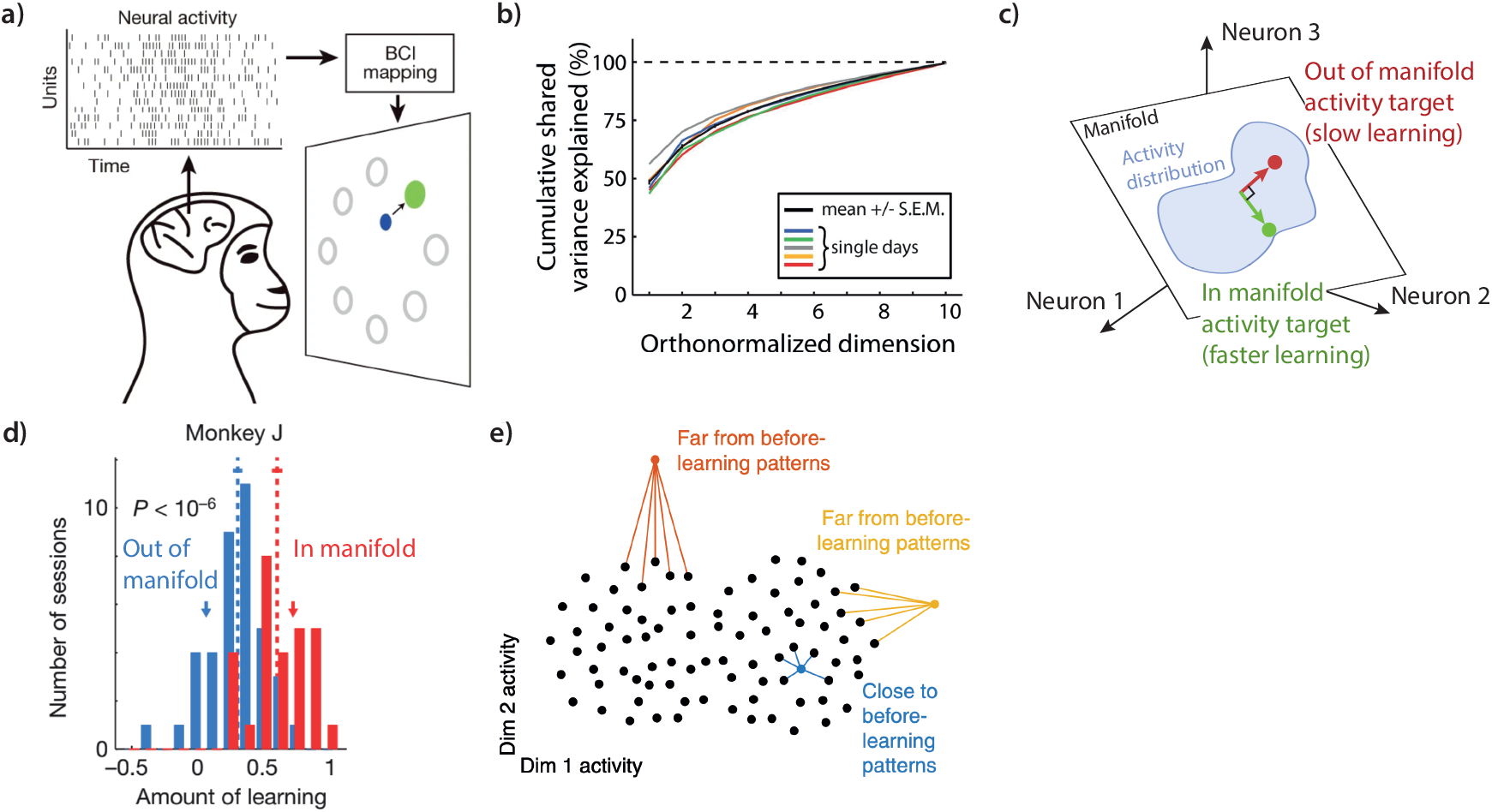
a) In a brain-computer interface (BCI) experiment, an animal must learn to control recorded neurons to solve a behavioural task. b) The neural activity measured by a BCI interface is typically much lower dimensional (as measured here by independent component analysis) than the number of recorded neurons (∼90 in this experiment). (a) and (b) are reproduced from Sadtler et al. (2014). c) Neural activity occurs within a reduced-dimensional manifold within the full space of allowable neural activity patterns (represented here as a 2D plane within a 3d volume). Significant differences in learning rate are measured for BCI tasks that are aligned with this manifold as compared to out-of-manifold tasks. d) Example data reproduced from Sadtler et al. (2014) showing differences in amount of learning for a Macaque learning in- vs out-of-manifold tasks. e) Further experiments have shown that, if the BCI task is modified, neural activity patterns remain within the same region of the neural manifold – new activity patterns remain close to before-learning patterns (blue). Reproduced from Golub et al. (2018).

We examine two key insights from recent BCI experiments. The first is the observation that the speed at which a BCI task can be learned is dependent on how closely the required neural activity aligns with pre-existing activity patterns. Experiments in macaques (Sadtler et al., 2014) have shown that learning “out-of-manifold” tasks is significantly slower than “in-manifold” tasks (Fig 1b-d).

A second observation from within-manifold experiments is that the overall set of neural activity patterns within the manifold does not change significantly over the course of training (Golub et al. (2018), Fig 1e), with “reassociation” between existing activity patterns and new outputs, but few or no new patterns observed outside of this space.

These experiments (Golub et al., 2018; Oby et al., 2019; Sadtler et al., 2014), as well as follow ups (Feulner and Clopath, 2021; Menendez and Latham, 2019; Wärnberg and Kumar, 2019), hypothesize (implicitly or explicitly) that these two phenomena reflect different specific constraints (such as specific patterns of neural connectivity, see Discussion for details), and are not general features of optimization algorithms (Hennig et al., 2021).

We trained artificial neural networks (ANNs) to perform simulated BCI tasks, in which the network activity at a hidden layer is read out as a simple analogue to a BCI recording device. We used these in-silico experiments and supporting theoretical analyses to study BCI learning dynamics under online gradient-based credit assignment algorithms. Our analyses suggest a simple explanation for both (1) in- vs out-of-manifold learning differences and (2) neural reassociation: these observations are a direct consequence of the dynamics of such gradient-based credit assignment algorithms. The only assumption required to explain the essential features of the empirical data is that the brain updates neural representations via a local estimate of the gradient of the task performance.

## Results

### Credit-assignment algorithms

BCI tasks require control of the feedback loop connecting a neuron’s activity to reward. How are learning signals that encode task performance translated into a synaptic weight update? Evidence from artificial networks suggests that changing activity deep within a neural network to optimise a learning signal is challenging without an estimate of causal effect (i.e. an estimate of the gradient of task performance with respect to neural activity). Effective use of such causal estimates will naturally lead to gradient-descent^1^ dynamics in the network updates.

In artificial networks, neural credit assignment is achieved using backpropagation-of-error (referred to as backprop) to propagate gradients from the network outputs to neurons. Backprop *computes* a local gradient using knowledge of the network structure. While backprop learning is widely regarded to be biologically implausible (Crick, 1989; Rumelhart et al., 1986), many speculate that the brain leverages backprop-like credit assignment (Bengio et al., 2015; Lillicrap et al., 2020; O’Reilly, 1996). In order to support BCI task learning, such an algorithm would also require a mechanism to determine that the (arbitrarily chosen) BCI output neurons represent the network outputs, and are therefore the correct neurons from which to propagate error information upstream.

While exact gradient backpropagation is likely to be biologically unfeasible, almost all hypothesised neural credit assignment algorithms use an estimate of a neuron’s causal effect. These algorithms lie along a spectrum trading complexity of implementation against accuracy of credit assignment. To gain insight into the learning dynamics of a BCI task, we consider algorithms at both ends of this spectrum (Fig. 2). The first algorithm, backpropagation, is precise but complex. The second algorithm, node (activity) perturbation, represents the other extreme, as it is a simple and local algorithm but represents an inefficient use of task information. In this algorithm, small perturbations are applied to the activity of each neuron during the forward pass through the network. These perturbations are then correlated with changes in performance in order to generate a learning signal. This algorithm only requires the task error to be broadcast globally through the network - each neuron can locally compute updates using information about itself, its inputs, and this global error (Seung, 2003; Werfel et al., 2005; Williams, 1992). Despite its biological plausibility, node-perturbation has not been found to scale beyond small-scale networks and simple tasks (Lillicrap et al., 2016; Payeur et al., 2021). Nevertheless, node perturbation has been found to be consistent with measured activity changes in BCI tasks (Legenstein et al., 2010).

**Figure 2.**
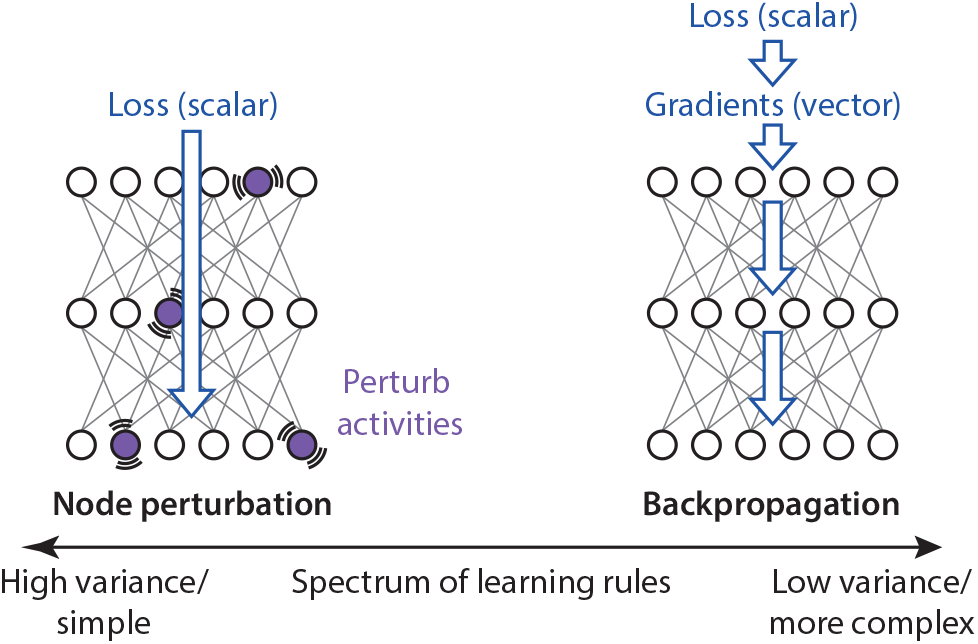
We study two learning rules, node perturbation and backpropagation, from a spectrum of rules that balance credit-assignment effectiveness against complexity. Backpropagation uses the chain rule to propagate gradients backwards through the network from output neurons. This requires potentially biologically-implausible additional network infrastructure but gives low-variance gradient estimates. Node perturbation is much simpler. In this algorithm, small perturbations are applied to the activity of each neuron during the forward pass through the network. A noisy estimate of the local gradient can be obtained by correlating the applied noise with changes to the overall performance. This algorithm only requires the scalar loss to be broadcast through the network.

As we will show, our theory and experiments support the claim that gradient-based (or hill-climbing) algorithms throughout the spectrum of precision of gradient updates display the same fundamental phenomena during BCI learning.

### In-silico BCI neural network models capture in- vs out-of-manifold learning differences

We run in-silico experiments using a 3 hidden-layer artificial neural network. This network is fully connected and purely feed-forward, yet even this simple model exhibits the phenomena of interest to us. Full details of our network implementation are given in Methods.

We first pre-train the network, using online gradient-descent with gradients estimated by back-propagation or node perturbation. We pre-train on an objective designed to be reminiscent of a straightforward behavioural task – a noisy image of a dot in one of ten positions arranged in a circle is input to the network, and the network must learn to output the two-dimensional coordinates corresponding to the dot position (Fig 3a). For us, this step is in loose analogy to the background learning that goes on during an animals normal lived experience. As we will see, learning a behavioural task naturally establishes structure in a network, resulting in in- and out-of-manifold patterns.

**Figure 3.**
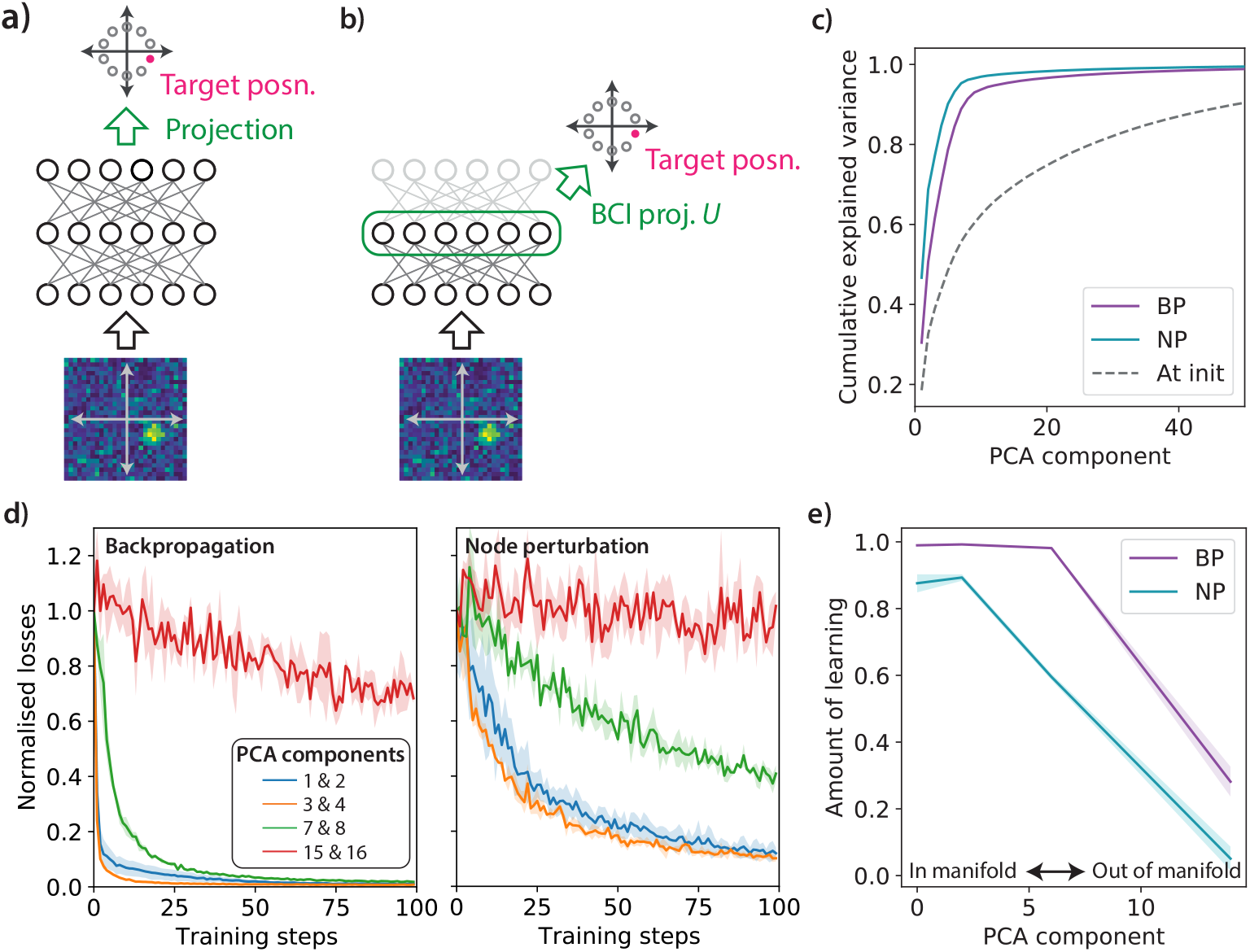
Model of a BCI experiment. a) A feed-forward network is pre-trained to map a noisy input image of a dot in one of ten possible positions to corresponding two-dimensional output coordinates (in our actual experiments three hidden layers of 256 units are used). b) To simulate the BCI task, the network activity at the penultimate layer is projected down to two components using a projection matrix *U* and read out. The network must again produce two-dimensional output coordinates corresponding to each of the ten positions, but now via *U*. c) As is observed in biological networks, most of the neural activity at the BCI layer is concentrated in a subset of the principal components for both backprop (BP) and node perturbation (NP), in contrast to the distribution of a randomly initialised network. d) Learning dynamics for BP and NP in experiments in which cursor coordinates are derived using a projection matrix *U* onto a specified pair of principal components. Learning is fastest for projections onto the largest components of the network activity. Shaded region represents standard deviation measured across 3 seeds. e) Total amount of learning (1 - normalised losses) for experiments in (d) after 100 training steps as a function of the projection principal component (specifically, the first component of the pair). As can be seen, more out-of-manifold tasks lead to less learning after a fixed number of training steps, in accordance with biological observations (Sadtler et al., 2014).

After pre-training, we model an intervention that is analogous to a BCI experiment, by instead utilising an objective based on the activity of neurons at the penultimate network layer^2^. This is achieved by linearly projecting this activity onto two-dimensional “cursor coordinates” using a fixed projection matrix *U*. This output is equivalent to the BCI cursor velocity in Sadtler et al. (2014), although we use the converted neural activity to directly represent the cursor position. The network must again produce a matching cursor position for each of the 10 different dot positions represented in the noisy input images (Fig. 3b), but via the BCI matrix *U*.

Principal component analysis (PCA) of the network activity at the BCI layer after pre-training reveals a similar spectrum to the macaque experiments (reproduced in Fig. 1b), with 90% of the neural activity variance captured by approximately 8 components, confirming the presence of an activity manifold (Fig. 3c). In real networks this manifold may be imposed by the dimensionality of the task (Gao et al., 2017) or neural connectivity (Wärnberg and Kumar, 2019). The specific mechanism(s) inducing this neural manifold do not impact our analysis or conclusions.

To investigate in- vs out-of-manifold learning dynamics, we set the BCI projection matrix *U* to be given by projections onto specified pairs of principal components. So for example, if we choose principal components 3 and 4, the network has to learn to modulate the 3rd (4th) principal component of its activity in order to control the 1st (2nd) dimension of the cursor position. We choose this slightly different mapping to those used in Sadtler et al. (2014), in which tasks are only labelled as either in- or out-of-manifold, in order to more clearly demonstrate the gradual changes in learning dynamics as tasks become increasingly out-of-manifold (higher principal component numbers).

As is shown in Fig. 3d & e, both backpropagation and node perturbation induce the same trend as was reported for biological experiments, with learning getting progressively slower for principal components that capture less of the neural activity variance (more off-manifold). In these feed-forward networks, there is no architectural constraint enforcing this. It is simply a natural property of gradient-based credit assignment algorithms operating on a network with structure induced by previous learning, as we will show theoretically.

### Theoretical analysis

So far we have provided empirical evidence that in-vs out-of-manifold learning differences are observed for gradient-based credit assignment algorithms in deep neural networks. We investigate this phenomenon theoretically by considering the learning dynamics of a simplified model consisting of a single-layer linear network *y* = *Wx* trained by online gradient descent with back-propagation or node perturbation, as shown in Fig. 4a (following Saxe et al. (2013); Werfel et al. (2005)). We think of this network as outputting the collection of neural activities recorded during a BCI experiment.

**Figure 4.**
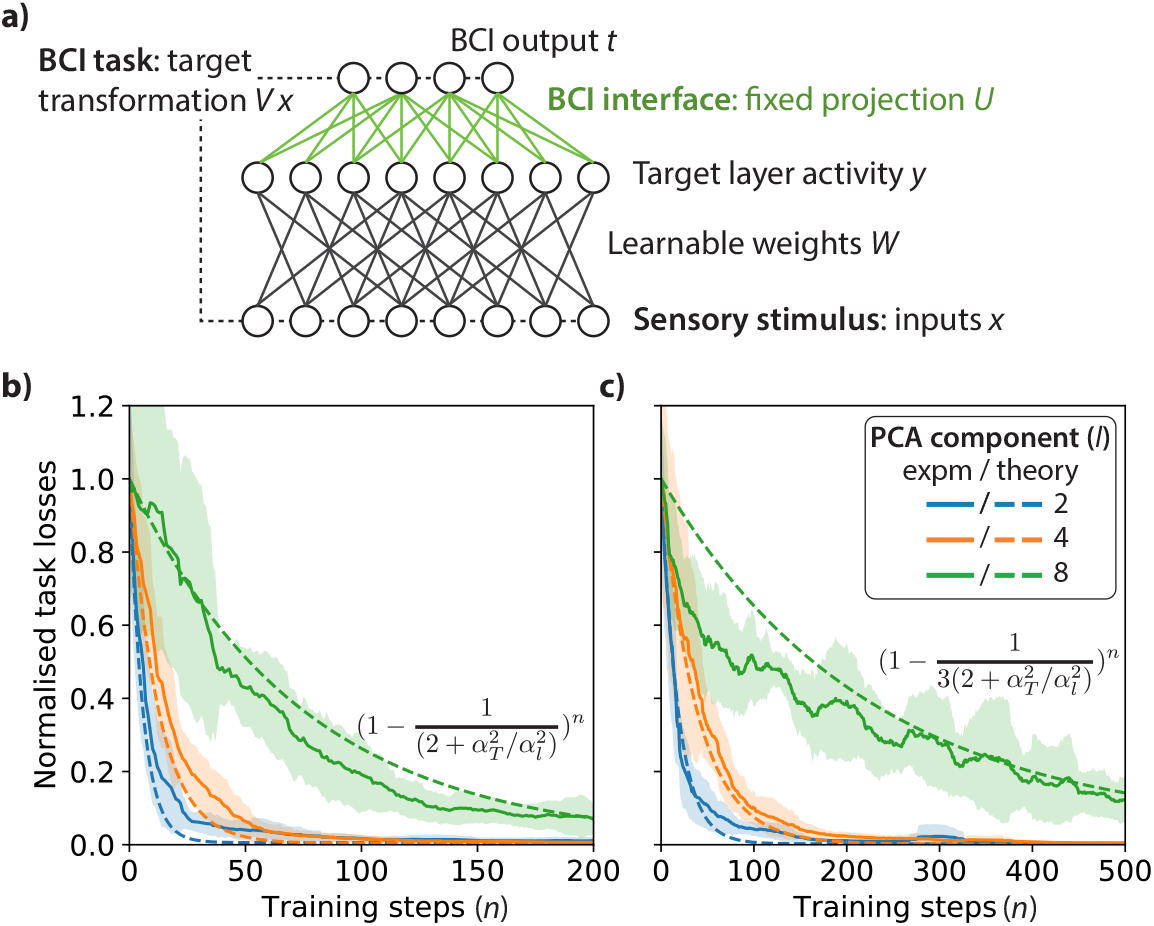
a) Simplified linear network model used to investigate in- vs out-of-manifold learning. Our theoretical analysis allows us to find approximate analytical expressions for the learning dynamics under certain restricted conditions (described in Appendix 1) – in particular, our theoretical analysis is of a 1D equivalent of the cursor task. Plotted are dynamics for (b) backpropagation and (c) node perturbation for BCI projections *U* onto varying principal component numbers (legend). As can be seen our theoretical predictions (dashed) closely match the learning dynamics (solid). This analysis confirms that as the target output principal component number increases (associated with smaller variances), the learning rate is suppressed.

To model the BCI task, we project the layer outputs *y* to a set of target dimensions *t* using a fixed projection matrix *U*, such that *t* = *U y* = *UWx*. Crucially, in contrast to previous related analyses (Saxe et al., 2013; Werfel et al., 2005), we explicitly consider the case where each component of *x* is not independent and identically distributed, but instead these components are derived from an underlying set of principal components *l* with potentially different associated variances 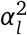. Taking these input principal component variances into account leads to in- and out-of-manifold learning dynamics for both back-propagation and node perturbation (Fig. 4b & c). In particular, we prove that the maximum achievable learning rate is determined by the ratio of the variance 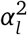 of the principal components that the BCI is targeting as compared to the total input variance 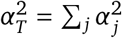. Full details of our analysis are given in Appendix 1. Our in- vs out-of-manifold learning results can be understood in intuitive terms: a BCI experiment requires an animal to produce specific activity patterns conditioned on specific inputs. If the target activity pattern is only a small component of the activity (out-of-manifold), it will be harder to pick the learning signal associated with it out of the variance induced by other components, and learning will be slower.

### Relation to biological networks

Biological neural networks are clearly not fully-connected feed-forward networks, and in particular contain significant recurrent connectivity. These additional details of biological networks may help explain the origin of the observed manifold of activities. However, our analysis is agnostic to this - what is essential is that there is a manifold of activity, as opposed to its precise underlying causes.

The essential results of our analysis translate to networks with recurrence and sparse connectivity. Recurrence changes the inputs to a given neuron, which are no longer simply determined by external sensory observations. In addition, this recurrence means that the consequences of perturbations must be measured over an extended time duration. However, it is still perfectly possible to measure the principal components of the neural activity, and all other relevant details should carry over to allow an algorithm such as node perturbation to operate with minimal modifications. Further, modelling suggests that feed-forward networks effectively capture the initial volley of activity in recurrent motor control circuits (Lillicrap and Scott, 2013).

Similarly, the structure of neural connectivity in biological networks will likely constrain the space of expressible neural activities. Nonetheless, for tasks such as the BCI experiments considered (Sadtler et al., 2014), the dimensionality of the neural activity is strongly determined by the input and task dimensionality (Gao et al., 2017) and is lower than would be expected from connection sparsity alone. This suggests that sparsity is unlikely to be the dominant factor in enforcing the observed manifold. The observation that out-of-manifold tasks can be learned over several days (Oby et al., 2019) shows that biological networks can support activity patterns outside of those initially observed, and hence further indicates that sparsity or other structural mechanisms do not restrict learning to be only in-manifold.

### Investigating changes in network activity distributions over the course of training

After showing that gradient-based credit assignment algorithms are consistent with in- vs out-of-manifold learning differences, we now turn to a related observation from recent BCI experiments. In these experiments (Golub et al., 2018), macaques were trained to solve a cursor-based BCI task as described previously. This task was then “perturbed” by substituting a new mapping connecting neural activity to BCI output positions. This perturbation led to an initial decrease in performance, but the monkeys were eventually able to recover, given sufficient trials. A key finding of these experiments was that the overall repertoire of neural activity patterns within the neural manifold did not change significantly over the course of training, although patterns within this repertoire may become associated with different targets.

We model this experiment in-silico. We first train a feed-forward network on a BCI task using backpropagation as described above, but with a randomly chosen projection matrix *U* onto the top 10 principal components. We then perturb the task by substituting a new matrix *U* connecting these PCA components to the cursor position (Fig. 5a). In keeping with the original experiments, we choose this matrix such that the initial perturbed outputs are close to the original outputs (see Methods).

**Figure 5.**
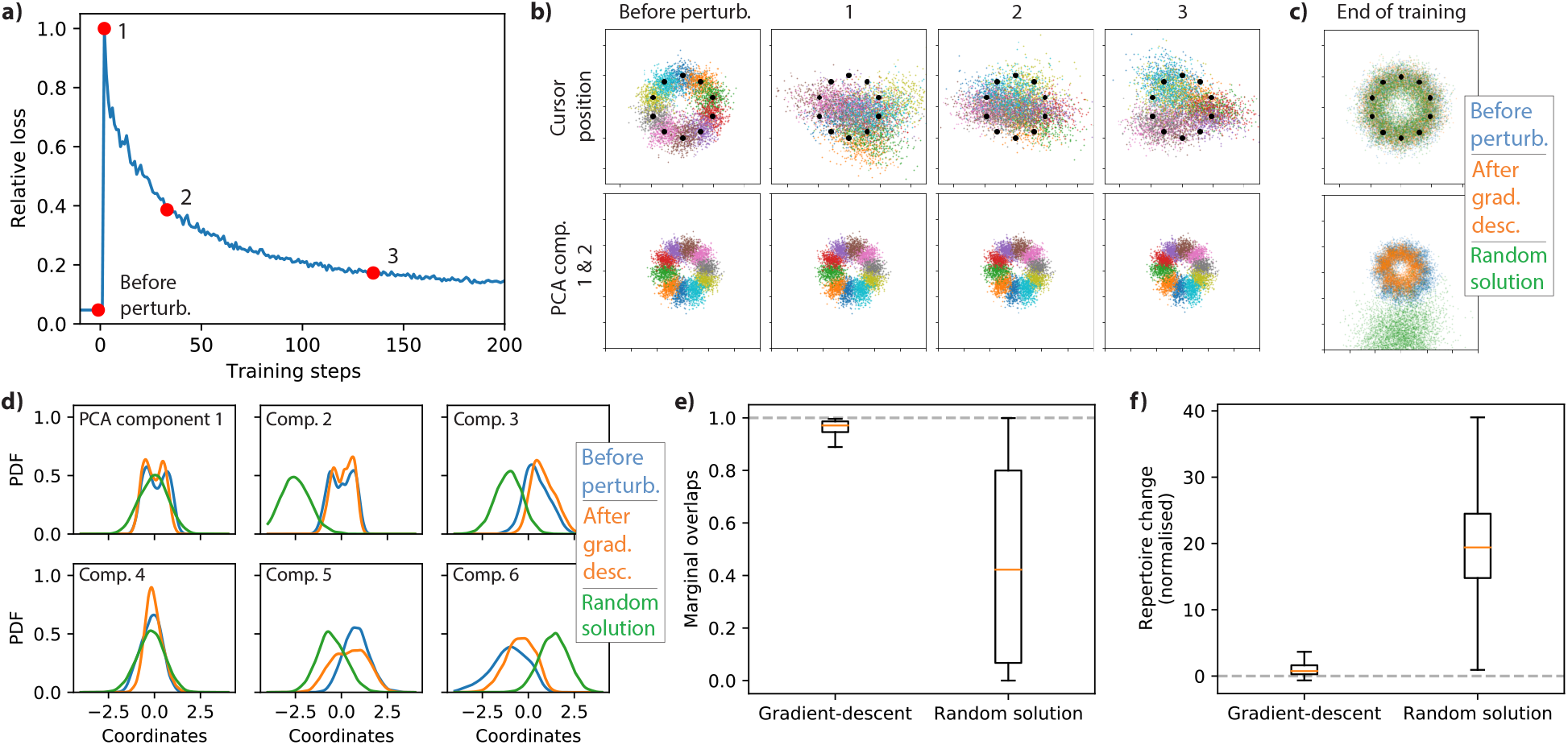
Changes in neural activity distribution are investigated by first training a network using backpropagation to solve a BCI task, and then “perturbing” this task by substituting a new matrix *U* connecting neural activity to cursor output positions. a) Training curves show an initial increase in loss after perturbation, followed by recovery over time. b) Visualisation of network activity changes over the course of training. Plots correspond to the positions marked by red dots in (a). The top panel shows output cursor positions (with black dots denoting the target positions), while the bottom panels show neural activity 1st and 2nd PCA components. Data is color coded by target. As can be seen, the distribution of cursor positions change significantly after the perturbation and then slowly returns to its initial configuration. In contrast, the PCA components are comparatively unchanged, as was observed in the original “reassociation experiments”. c) Overlaid distributions from (b) of cursor positions and PCA components before the perturbation (blue) and after training by gradient descent (orange). Further plotted are distributions for an example randomly initialised network trained on the same task (green), which converges to a very different distribution in PCA component space while outputting a similar cursor distribution. This qualitatively demonstrates that solutions found by gradient descent are much closer to the initial distribution than networks randomly chosen from the space of valid solutions. d) Example marginal distribution probability density functions (PDF) of neural activities along the top 6 principal component directions before the perturbation (blue) and after (orange) gradient descent for the experiment shown in (a) and (b), along with marginals for the randomly initialised network trained on the same task (green). e) Box and whisker plot of the overlaps between the before-perturbation and post-training marginal distributions for the data shown in (d) (but calculated for the top 10 PCA components and from 10 different experiments). This quantifies the qualitative observation in (c) – random solutions have lower marginal overlaps than gradient descent training. e) Similarly, following Golub et al. (2018), instead of examining marginals, we can measure distances between activity patterns in the full space of the top 10 principal components, and use this to calculate a “repertoire change” (see Methods), in which no change in distribution would give a near-zero repertoire change. Gradient-descent solutions induce such a near-zero repertoire change, in accordance with biological experiments.

Fig. 5b shows the evolution of the network outputs and the 1st and 2nd PCA components of activity over the course of post-perturbation training. As can be seen, the cursor positions are initially distorted by the perturbation, but eventually return to their original configuration. In contrast, the PCA components are comparatively unchanged by this perturbation and subsequent training. This is further visualised in Fig. 5c, in which the overall distributions of initial and final cursor positions and initial and final principal components are overlaid.

As the output of the BCI matrix *U* is much lower dimensional than the number of recorded neurons, there are many possible neural repertoires that would be compatible with the target BCI signal. This is a point that is explored in Golub et al. (2018) – many different transformation schemes for the neural repertoire, such as a smooth rescaling of activities, could all in principle support BCI task learning. To compare the solutions found by gradient-descent to the broader set of possible neural repertoires that can solve this task, we randomly initialise networks and train them to solve the same task (i.e. with the same matrix *U*). An example resulting repertoire is plotted in green in Fig. 5c – it clearly does not overlap as closely with the initial neural repertoire as the solution found by gradient descent. As we explore more quantitatively below, these experiments show that gradient-based credit assignment algorithms find solutions that are closer to the initial distribution than repertoires drawn randomly from the full space of possible solutions. Neurons are thus “reassociated” with downstream targets under the perturbation.

In Fig. 5e & f we make quantitative comparisons of the activity distributions for a set of 10 in-silico experiments run as described above, comparing this to the activity distributions for randomly initialised networks trained on the same tasks. As a simple metric, we examine the overlaps (Bhattacharyya distance) of the marginal distributions of the top 10 PCA components before and after gradient-descent training (Fig. 5d & e, see Methods). In addition, following Golub et al. (2018), we quantify the degree of change in the full (non-marginalised) space of the top 10 PCA components by calculating a normalised “repertoire change” (Fig. 5f, see Methods). These measurements further demonstrate that the distribution under gradient-based algorithms remains much closer to the initial distribution than would be expected based on randomly sampling from the space of valid network solutions.

These results can be understood intuitively. Gradient-based algorithms operating in overparameterized regimes naturally tend to find solutions close to the original space of network activities, and thus are consistent with “reassociation”. For an under-constrained problem such as a BCI task, neural activities will stay much closer than would be expected for solutions randomly sampled from across the space of valid parameters.

### Inducing out-of-distribution network activity patterns

A prediction of our work is that it is possible to induce more significant changes away from the initial distribution by modifying task structure^3^. In Fig. 6 we show similar experiments, but for which, in contrast to the initial experiments and the biological experiments presented in Golub et al. (2018), we no longer constrain our perturbation *U* matrix to ensure that the initial perturbed outputs are kept close to the original distribution. In addition, we reduce the number of PCA components that the BCI matrix *U* projects on to. This means that there is less degeneracy in the space of PCA component activities that can solve the task to a given level, and so greater differences in final versus initial distributions in the PCA component space are observed.

**Figure 6.**
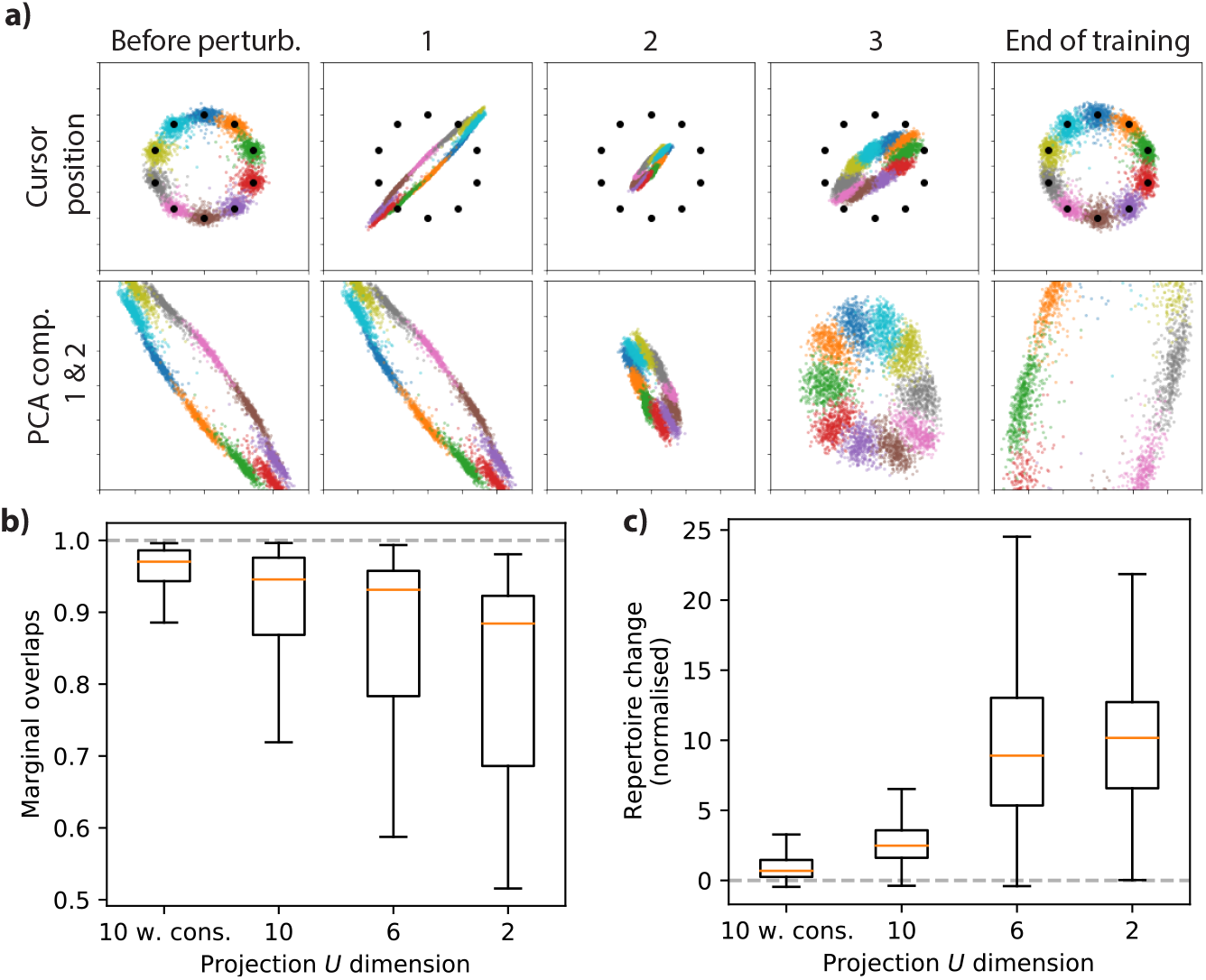
a) An equivalent perturbation experiment to that shown in Fig. 5, but with the cursor position derived from the top 2 principal components, instead of the top 10. In addition, we do not constrain *U* to ensure that the initial perturbed outputs are kept close to the original distribution. Plots labelled 1,2 & 3 correspond to measurements made at the same relative loss values as in Fig 5. Qualitatively, greater changes are visible in the 1st and 2nd principal components of the activity. b) Overlaps of the marginal distributions and c) normalised repertoire change for perturbation experiments with output projections *U* from the top 2, 6 and 10 PCA components, along with the equivalent 10D projection with *U* constraint results from Fig 5e & f (“10 w. cons.”). As can be seen, these less under-constrained projections induce larger shifts in the activity distribution.

## Discussion

Brain-computer interfaces hold promise for functional restoration and perhaps even functional augmentation. They also provide a powerful tool for fundamental neuroscience research, allowing unique control over the connection between neural activity, external feedback, and reward. This makes them ideal probes for investigating the algorithms used by the brain to learn behavioural tasks.

Hennig et al. (2021) have argued for an optimization view in understanding how learning unfolds in the brain, but posit that “key features of how neural population activity changes throughout learning that cannot be readily explained in terms of optimization and are not typically features of ANNs”, including speed of learning for in- vs out-of-manifold tasks. We instead find that differences in learning for in-versus out-of-manifold tasks are indeed a feature of typical ANNs trained by standard methods. Indeed both this in- vs out-of-mainfold difference, along with experimentally-observed neural reassociation, can be explained by gradient-based credit assignment mechanisms at both extremes of simplicity versus sophistication: backpropagation and node perturbation. This shows that these learning dynamics can be induced by a broad class of credit assignment algorithms in between these two extremes. This lends weight to an even stronger optimization view than that proposed in Hennig et al. (2021) – in fact behavioural/task optimization is sufficient by itself to explain many of the phenomena that they identify, without any further assumptions. It is possible that these phenomena will ultimately be explained by other aspects of the brain. However, we argue that, by Occam’s Razor, this simplest explanation should be preferred.

Gradient-based (hill-climbing) algorithms are likely necessary in the brain in order to explain findings such as the possibility of BCI control in the first place. Nonetheless, it is worth emphasising that not all hypothesised neural update mechanisms are gradient-based. Two-factor Hebbian (Hebb, 1949) or BCM (Bienenstock et al., 1982) learning are error-signal insensitive and therefore do not follow a task gradient. Rather, applied on their own, such updates will tend to move in a random direction relative to the task. On the other hand, various random search algorithms (such as simulated annealing and some genetic algorithms) will improve task performance over time, but operate by non-local changes to synaptic parameters or the associated neural activity/representations. This is contradicted by experimental data. Finally, it could be that neurons are optimising a different objective than the behavoural task in question, or may be optimising more than one objective function (Hennig et al., 2021).

Previous studies have explored different possible explanations for in- vs out-of-manifold learning differences (Feulner and Clopath, 2021; Hennig et al., 2021; Menendez and Latham, 2019; Wärnberg and Kumar, 2019). Wärnberg and Kumar (2019) suggest that local connectivity patterns constrain activity to a low-dimensional manifold. They argue that, for such networks, inducing out-of-manifold activity will require larger weight changes. They propose that these larger weight changes will require more extensive learning and hence will occur more slowly. Central to their paper is the hypothesis that the manifold dimension in biological neural networks is determined primarily by network structure. However, there is evidence that this is not the case, with the dimensionality of recorded activity also determined by the task complexity (Gao et al., 2017). The two key assumptions made by Wärnberg and Kumar (2019), that manifold dimensionality is determined primarily by connectivity, and that larger weight changes are intrinsically slower to learn and dominate learning rates, are not required in our analysis.

Feulner and Clopath (2021) study BCI task learning in a recurrently connected network, with a similar simulated BCI readout to our experiments. They find that final performance for in- vs out-of-manifold learning is the same under backpropagation, and therefore posit that backpropagation is not sufficient to explain in- vs out-of-manifold learning. However, as is shown in their Supp. Fig. 6a, the learning rate for out-of-manifold learning is slower. This result – slower learning, but convergence to the same eventual performance, is in line with our theory and experiments. Their difference in interpretation seems to be because they focus on final performance, which we argue is not the relevant metric. As is shown in Oby et al. (2019), animals are eventually able to learn out-of-manifold tasks, just at a slower rate than for in-manifold tasks.

Summarising previous works, we acknowledge that there are other mechanisms that could explain differences in in- vs out-of-manifold learning. However, we propose that none of these mechanisms are necessary on top of the assumption of gradient-based credit assignment algorithms. It is likely that virtually all such hill-climbing learning rules directly induce these dynamics, and it is hard to reconcile the very existence of BCI task learning without assuming that the brain carries out some form of gradient-based credit assignment. The universality of these dynamics is underlined by the fact that they are observable in simple feed-forward fully connected networks.

Re-association is another phenomenon that we argue is well explained by an optimization-based view of learning. Both artificial and biological neural networks tend to be over-parameterised, and experimental BCI objectives are typically under-constrained. In this context, gradient-based dynamics naturally lead to solutions that are close to the original weights and activity distributions. This is simply because there tend to exist many solutions nearby in parameter space that satisfy the objective (Dauphin et al., 2014). Hill-climbing updates that use local gradient estimates will thus tend to discover solutions nearby in parameter space. Thus, we shouldn’t be surprised to see re-association as a phenomena in the brain. Rather, we should be surprised when we do not see it. This idea is echoed in recent machine learning literature, which has shown that it is possible to quickly adapt large pre-trained networks to a broad range of downstream tasks of interest via “fine-tuning” paradigms (Brown et al., 2020; Radford et al., 2019; Reid et al., 2022). As the name suggests, fine-tuning induces only small changes in the network representations (Zhou and Srikumar, 2021), suggesting that representations in the network can be quickly “re-associated” with new functionality for the downstream tasks.

Our theory and experiments allow us to explain previously perplexing learning phenomena using only a simple conceptual model. Instantiations of this model also let us make empirical predictions that could be tested in BCI experiments in the laboratory. Fig. 6a illustrates one such prediction, wherein we expect to see that BCI tasks with *less redundancy* will lead to greater changes in neural activity patterns. In this example, mapping only two principal components to cursor positions should induce greater changes (as observed within the principal component manifold) than using ten principal components. Our in-silico experimental paradigm can be used to derive many more predictions.

Given our conclusion that BCI experiments are consistent with gradient-based learning, we believe that a key area for future research will be to narrow down the specific algorithms supporting credit assignment. Specifically, we believe that the following questions are critical:

- Where is the plasticity occurring that underlies observed activity changes? This may be at the recording site or in a brain region upstream of the BCI target region.
- What kinds of reward signals are transmitted to the recorded brain region? (e.g. a vector from other tissue, or a global scalar signal)
- Is the brain using any internal notions of it’s own function (e.g. gradient estimates) that it uses to do credit assignment?

Many of these questions cannot be resolved using BCI alone. For example, BCI can reveal activity changes, but does not directly probe underlying connectivity changes. In addition, further techniques will be needed to track the delivery of dopamine/reward related signals into cortex over the course of learning.

It may not be straightforward to make algorithmic inferences based on observations of a neural system that is learning. A handful of studies have begun to concretely ask how to perform these inferences (Nayebi et al., 2020; Portes et al., 2022). For example, Nayebi et al. (2020) show that it is possible to distinguish between standard backprop and feedback-alignment (Lillicrap et al., 2016) learning based on activity updates, however, this relies on having known examples in-silico to train classifiers. Portes et al. (2022) investigates how a biased update derived from a credit-assignment model can be distinguished from an unbiased reinforcement learning algorithm using BCI experiments. It’s an exciting research project to build on these investigations to find evidence for specific biological network update rules.

In closing, we return to the key motivation that, beyond supporting novel neuroscientific research, BCI interfaces show promise for improving the quality of life for patients with a range of disabilities. A relevant question is therefore how to translate insights from BCI research into improved BCI interfaces. Typically the neurons used for a BCI interface are chosen based on practical constraints, but there is significant freedom in how the raw recordings from these neurons are translated into control signals. A better understanding of the learning dynamics and representational changes underlying BCI learning should help better guide these design choices. For example, it is clear that fast learning of BCI control can be achieved by ensuring that desired changes align with the observed neural manifold. Given that previous experiments have suggested that this manifold is not innate, but is determined by the task statistics, it may be possible to use carefully chosen inputs or even neural stimulation to shape the input manifold, and therefore increase the variance within the target dimensions in order to boost learning.

## Acknowledgements

We thank Márton Rózsa, Gregory Wayne, Claudia Clopath and Agnieszka Grabska-Barwinska for discussions.

## Author Contributions

PCH carried out the investigations and analysis. KD and KS provided domain expertise. PCH and TPL wrote the paper with feedback and guidance from KD, KS and MB.

## Methods

We used the simplest possible in-silico setup to probe the questions of interest. All in-silico experiments were implemented using Python & JAX (Bradbury et al., 2018). We used feed-forward networks with 3 hidden layers of 256 units and an output layer of 10 units. The neurons at the hidden layers used a leaky-ReLU activation function, apart from the final hidden layer, which had an identity activation in order to simplify the implementation of the “BCI” experiments.

We first pre-trained the network on noisy input images of a dot at one of ten positions on the perimeter of a circle. The corresponding target cursor coordinates consisted of ten points uniformly arranged on the perimeter of a circle centred at (0, 0) and with 0.5 radius. The network was trained to output these targets using a standard mean-squared error loss and a stochastic gradient descent (SGD) optimiser. We note that the precise pre-training objective utilised is not critical in our results - we have observed the same phenomena with several such objectives.

Next, in order to simulate a “BCI” experiment, we instead “recorded” the penultimate network layer as the network “output”, and projected this layer’s activity to two dimensions using a fixed projection matrix *U*. We trained this network output using the same task as for pre-training, but now using these different network outputs.

Backpropagation was straightforward to implement using standard JAX functionality. We used custom code to generate the node perturbation gradient estimates. This was implemented in a simple manner by first running the network forwards with no perturbations, and then running it again forwards with the same input, but with perturbations applied to each neuron in the network (Werfel et al., 2005).

For all experiments we fixed the batch size to 1000 and swept the optimiser learning rate *η*, choosing the learning rate that led to the best performance after a fixed number of training steps. Note that there is no distinction between a training and validation set in our experiments because the training points are generated online.

### Reassociation experiments

For our reassociation experiments, we first trained a network as described above, but with a fixed random projection matrix *U* onto the top 10 principal components of the final hidden layer activity. This matrix was generated by randomly sampling each component from a standard normal distribution scaled by 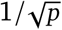 (to preserve variance), where *p* gives the number of principal components projected onto (here 10).

We then chose a new random matrix *U*, as a perturbation roughly equivalent to those investigated in Golub et al. (2018). In order to match these reference experiments, we chose this perturbation matrix such that the resulting initial network performance was not too dissimilar to the unperturbed performance. Specifically, we rejected perturbations for which the standard deviation of either cursor coordinate over a sample of 5,000 neural activity patterns was more than 4% different to the initial standard deviation, and for which the either cursor coordinate had an absolute mean value greater than 0.03. These filters were chosen somewhat arbitrarily, but aimed to have a net effect of accepting roughly 1 perturbation per 100,000.

### Reassociation experiments with different dimension projections

For reassociation experiments in which we adjusted the number of principal components *p* that the projection matrix *U* projected onto (Fig 6), we used a larger number of train steps (10,000), as the lower dimensionality projections occasionally took more steps to converge to a given loss level.

### Marginal overlaps

We calculate the marginal distributions of the principal components of the neural activity using standard kernel density estimation methods. The reported overlaps between these marginal distributions correspond to their Bhattacharyya distances. These are calculated by numerically integrating the square root of the product of their respective kernel density functions.

### Neural repertoire change

In order to aid comparison with the results from Golub et al. (2018), we adopt their “neural repertoire change” metric to compare distributions of neural activities.

We start with a series of *n*_*i*_ = 5000 measured activity patterns ***z***_*i*_ in a 10D principal component space, drawn from the initial distribution of network activities (before perturbation). Given these activities, we can calculate a corresponding covariance matrix ***S***_*i*_. Now, given a different series of *n*_*f*_ = 5000 measured activity patterns ***z***_*f*_ in the same space of principal components, but drawn from a different distribution, we can assess the distribution overlap by calculating a neural repertoire change as follows:

We first calculate the Mahalanobis distance *d*_*M*_ from each activity pattern 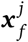 in ***z***_*f*_ to each activity pattern 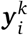 in ***z***_*i*_, given by:

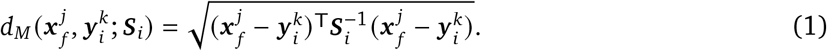

For each 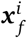 in ***z***_*f*_, we keep the corresponding first nearest-neighbor distance *p*_*i*_.

In order to normalise these distances, we assess the corresponding first nearest-neighbor distances between patterns in the original distribution in an equivalent way, by calculating 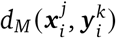 for all activity patterns 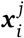 and 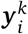 in the original distribution. We take the mean *ν* of the resulting first nearest-neighbor distances. The resulting normalised first nearest-neighbor distance for 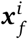 is given by:

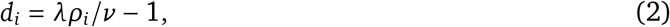

where λ = (*n*_*i*_ − 1) /*n*_*i*_. This is reported as “neural repertoire change” in Golub et al. (2018). This metric is calculated such that patterns drawn from the same distributions should show a change of near zero, while increasingly distinct distributions should show increasing repertoire changes.

## Appendix

### 1. Theoretical analysis of BCI experiment learning dynamics

Here we provide further details of our theoretical investigation of in- vs out-of-manifold learning differences in gradient-based credit assignment algorithms.

We consider a simplified model consisting of a single-layer linear network *y* = *Wx* (following a similar approach to Saxe et al. (2013); Werfel et al. (2005)). Here, *y* plays the role of the recorded neurons in physiology experiments. In order to model the BCI task, we project the outputs *y* to a set of target dimensions *t* using a fixed projection matrix *U*, so that *t* = *Uy* = *UWx* (Fig. 4a). The target for these outputs is defined as *t*_*T*_ = *Vx*, with a corresponding mean-squared error loss 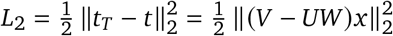.

### 2. Whitened and uncorrelated inputs

We follow similar derivations to those in Werfel et al. (2005) and study how the difference *V* between the target transformation *V* and the composite transformation *UW* evolves over training. Importantly, as in Saxe et al. (2013); Werfel et al. (2005), we begin by initially assuming uncorrelated and whitened inputs *x*. This assumption does not hold in typical BCI experiments (Fig 1), for which the dimensionality of the activity is significantly lower than the number of neurons. The extension to correlated inputs will be key to understanding in- vs out-of-manifold learning. However, we begin with uncorrelated inputs both for didactic clarity, and because they reveal interesting insights into the learning dynamics of the networks.

#### 2.1. Backpropagation

We first consider learning under online gradient descent (i.e. a batch size of 1) using gradients estimated by backpropagation.

Given an input *x*, the network loss will be 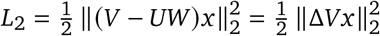. The corresponding gradient with respect to the weights will be

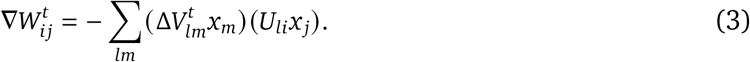

Applying a gradient descent update step on 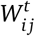 gives

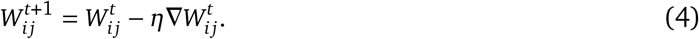

Instead of tracking changes in the network loss, it is more straightforward mathematically to consider changes in the transformation error 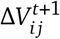:

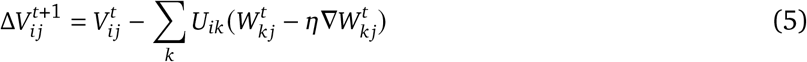

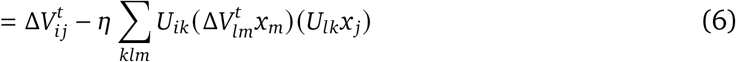

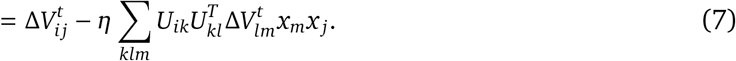

We square this to get an expression for the squared transformation errors, which we will subsequently relate to the loss *L*_2_. Taking the expectation of these squared errors over uncorrelated and whitened inputs *x* shows that 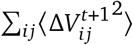 evolves under one step of gradient descent as:

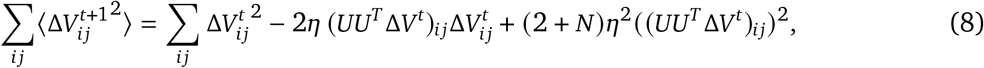

where *N* is the input dimensionality.

In the most general case this expression can lead to complex dynamics since the different elements of Δ*V* are coupled via the BCI projection matrix *U*. However, in the case where the projections *U* are mutually orthogonal such that 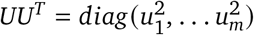, this reduces to:

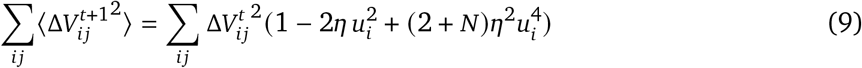

This means that target components *t*_*i*_ with different associated *u*_*i*_ will have different optimal learning rates 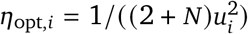. It is therefore important for optimal learning to ensure that *u*_*i*_ are the same for all of the target outputs.

For an orthonormal BCI projection matrix *U*, the above expression reduces to the equivalent expression in Werfel et al. (2005) for a single layer network:

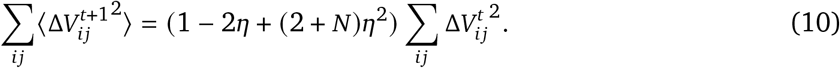

In the case of uncorrelated and whitened inputs, Δ*V* can be related to the expected loss by 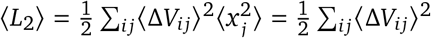. Therefore, we can find that (using the recursion relation given in Eqn. 10):

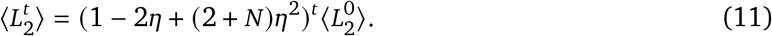

As can be seen, ⟨*L*_2_ ⟩ will decrease as long as (1 − 2 *η* + (2 + *N*) *η*^2^) *<* 1.

It is worth highlighting an important ramification of Eqn. 11 - the learning dynamics of backpropagation in such a network are insensitive to the number of neurons *M* at the output layer *y*, and are entirely determined by the input dimensionality *N*.

#### 2.2 Node perturbation

It is possible to derive similar expressions for node perturbation.

As for backpropagation, the network loss will be 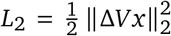. Node perturbation applies a perturbation drawn from a normal distribution *N*(*μ* = 0,*σ*) to each neuron in *y* and correlates this with the change in *L*_2_. Given this perturbation, an unbiased gradient estimator for Δ*y* is

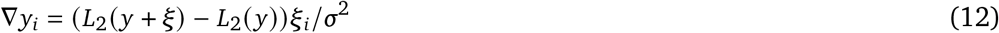

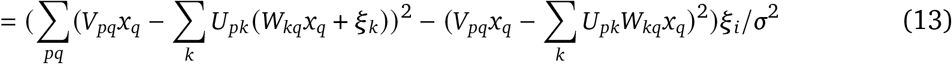

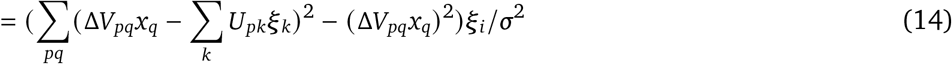

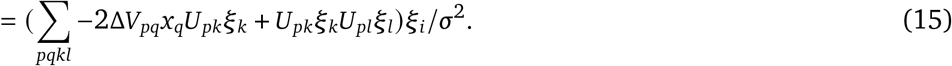

The corresponding weight gradients are ∇*W*_*ij*_ = ∇*y*_*i*_*x*_*j*_. This can be used to update each weight according to:

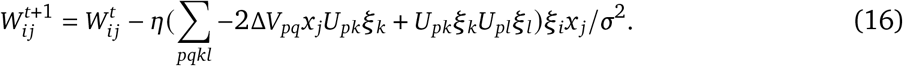

Following a similar line of reasoning as used above for backpropagation, for orthogonal *U*, this again reduces to the expression for a single layer network derived in Werfel et al. (2005):

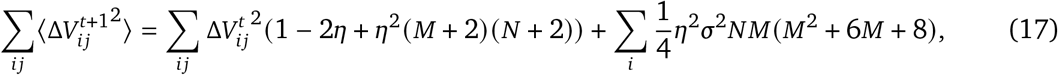

where *M* gives the target *t* dimensionality (not the number of neurons). As is the case for backpropagation, for orthogonal *U*, node perturbation is insensitive to the number of output neurons in *y*, depending only on the input dimensionality *N* and target dimensionality *M*.

### 3. Extending the analysis to correlated neural inputs

Our analysis so far has relied on the assumption of uncorrelated and whitened inputs (as in Saxe et al. (2013); Werfel et al. (2005)). For real data, or for neural inputs deep within a network, this is typically not the case. Removing these assumptions leads to further informative insights about the learning dynamics of BCI tasks. In order to make progress in studying this analytically, we have to transform the problem into a basis given by the input principal components.

Specifically, we assume that the neural inputs *x* can be expressed in terms of principal components *p*, such that *x* = *Pp* (Fig 7). These components will be uncorrelated, but not whitened, with a different variance 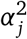 for each input principal component *p*_*j*_.

**Figure 7.**
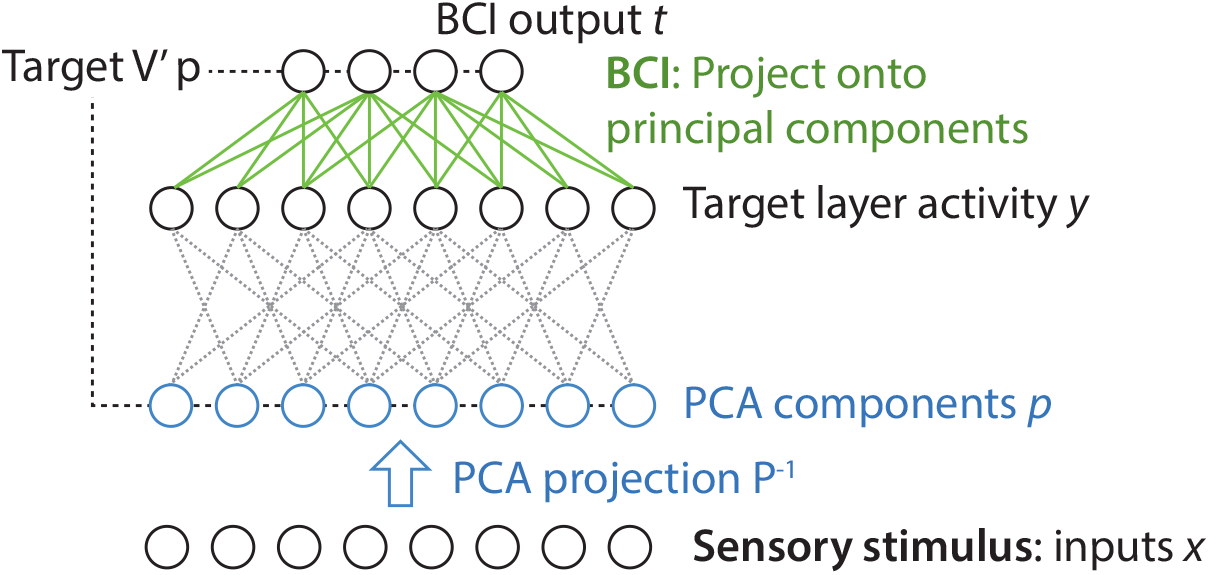
a) Linear model of a BCI experiment with correlated inputs. Inputs *x* are transformed into an uncorrelated basis *p* by projecting onto principal components. The BCI interface *U* is assumed to be a projection onto (a subset) of the principal components of the target layer *y*.

Analyzing our model network in this basis, we find that the total network transformation is *t* = *UWPp*. The corresponding network targets are *t*_*T*_ = *VPp*. Instead of studying the dynamics of Δ*V*, we can instead investigate Δ*V*^′^, = (*VP* − *UWP*).

For backpropagation, the learning dynamics of 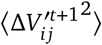 are given by

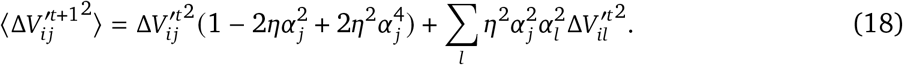

This shows that online gradient descent induces coupling between elements 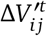 and 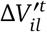 connecting different inputs *j* and *l* to the same output *i*, with a strength determined by 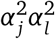. If one input *l* has a high variance 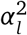, it can induce increases in the 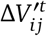 elements associated with other inputs *j*. The smaller variances 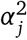 associated with these other inputs means that these induced errors will then only be corrected at a slower rate. This can be the case even if 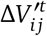 starts out at zero, as shown in Fig. 8.

**Figure 8.**
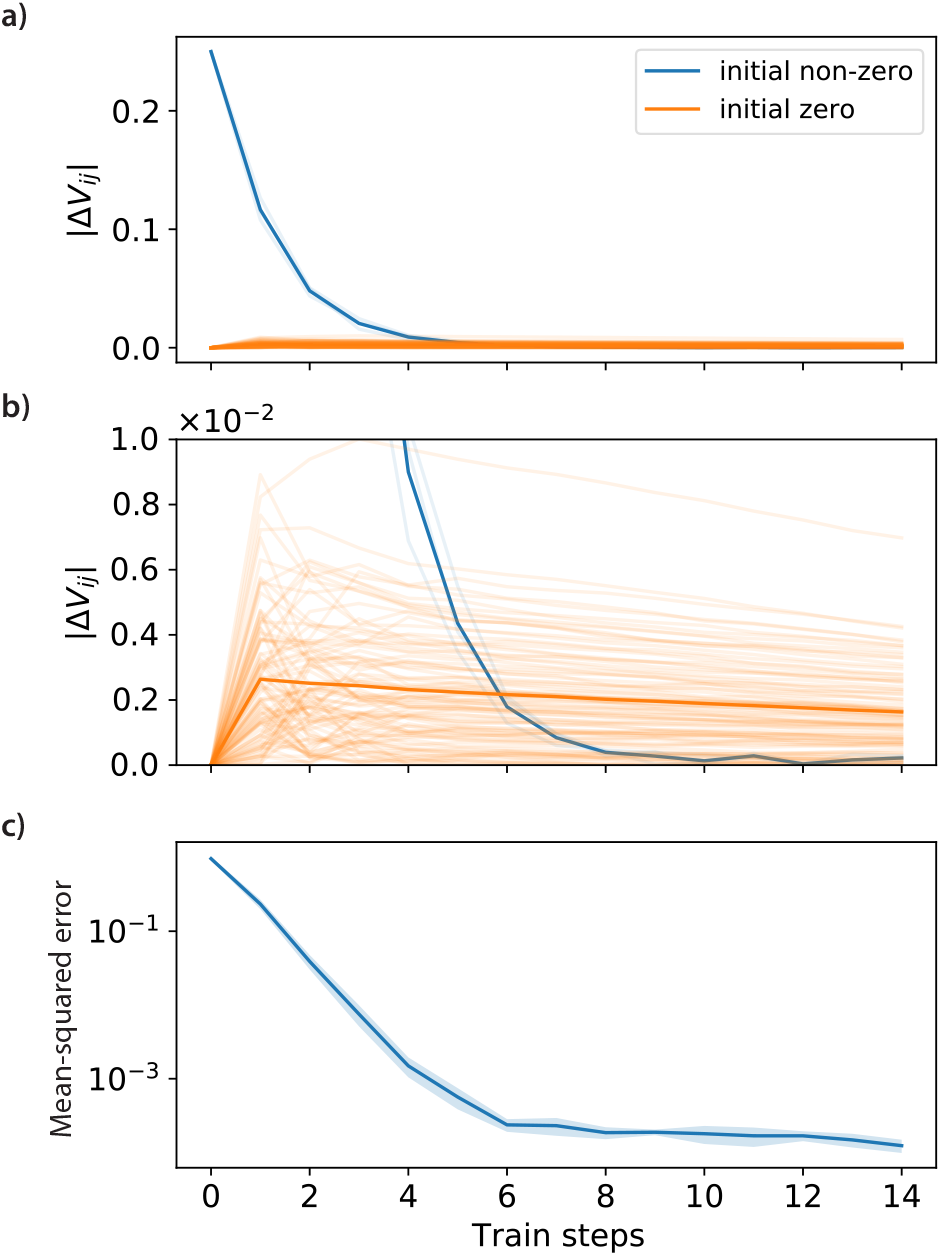
Didactic example of the learning dynamics for a neural network with a 32 dimensional input and a 1 dimensional output target. In order to clearly demonstrate the impact of spurious correlations, one input component has 16 times the variance of the other input components. The target transformation *V* is the zero matrix, and the only weight of *W* that is initially non-zero is that associated with the high-variance component (labelled “initial non-zero”). (a) As can be seen, this weight is rapidly suppressed, but at the same time increased weights are induced for the other initially zero components (labelled “initial zero”). The same data are shown in (b), but re-plotted to more clearly show the induced weights. c) Ultimately these induced errors dominate the residual loss.

Loosely put this means that, for a given desired input-output connection, the stronger the other inputs to the output are, the harder it is to follow the true learning signal as opposed to changes induced by spurious correlations between the inputs and the output.

#### 3.1 Node perturbation

A similar expression can be derived for node perturbation:

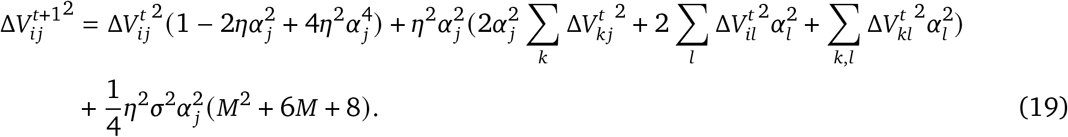

This has similar contributions to its dynamics, with elements associated with different principal components coupled according to their input variances.

### 4. In- and out-of-manifold learning with correlated inputs

In a setting such as BCI experiment, we do not have control over the distribution of input variances 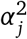. However, we do have control over *U*, the neural projection used to define the BCI task. In particular, we can choose whether *U* defines an “in-” or “out-of-manifold” projection from the output activities *y*.

A full analytical analysis of the learning dynamics in even this setting is intractable, but we can gain some insight by considering the simplest possible setting, in which only one component of the transformation error 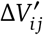 is initially non-zero (for *i* = *k, j* = *l*). This approximately corresponds to a BCI experiment in which a single-dimensional cursor output (*k*) must be controlled via a specified principal component (*l*) of the network activity. We hence take *M* = 1 in our analysis.

For an initial step of gradient descent under these conditions, it can be determined from Eqn. 18 that the minimum achievable value of 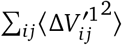 is given by:

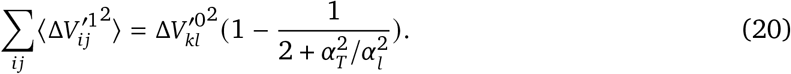

The decrease in error is determined by the ratio between the variance 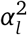 associated with the target component (*l*) and the total input variance 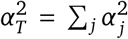. The greater this ratio is, the greater the decrease in 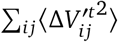 that is achievable under one step of gradient descent.

Eqn 20 is only valid for the first step of gradient descent in learning (as after the first step there will be further non-zero components of 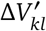). However, if we assume that the other components remain comparatively small, at least for the initial stages of learning, we can iteratively apply the expression in order to predict the evolution of 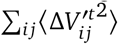 over time. This gives:

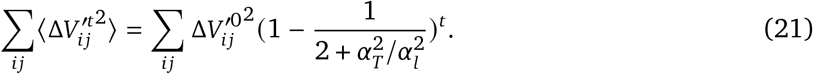

A similar analysis for node perturbation can be carried out based on Eqn 19. This gives an equivalent expression for the evolution of 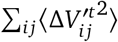 over time:

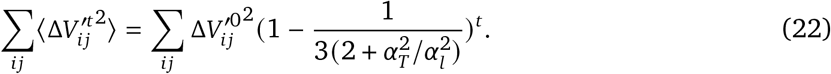

These equations give approximate expressions for the evolution of the weight errors 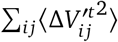. For inputs with varying principal component variances 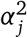, this is not directly proportional to the loss *L*_2_, since 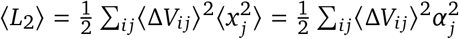. Nonetheless, if one element of the weight errors is dominant, we might expect that the *L*_2_ is approximately proportional to 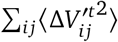.

In Fig. 9, we show the simulated learning dynamics for a network under the conditions outlined above, in which a single-dimensional cursor output *t* must be controlled via a single principal component of the network activity. We overlay our predictions for ⟨*L*_2_⟩ and 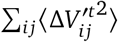 over time based on Eqns 21 and 22. We expect that these approximate expressions will only be valid for the initial stages of learning. We therefore match the equations to the data by assuming a fixed offset, determined by the values at the end of the training period. We additionally set the initial 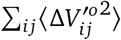 to the measured value.

**Figure 9.**
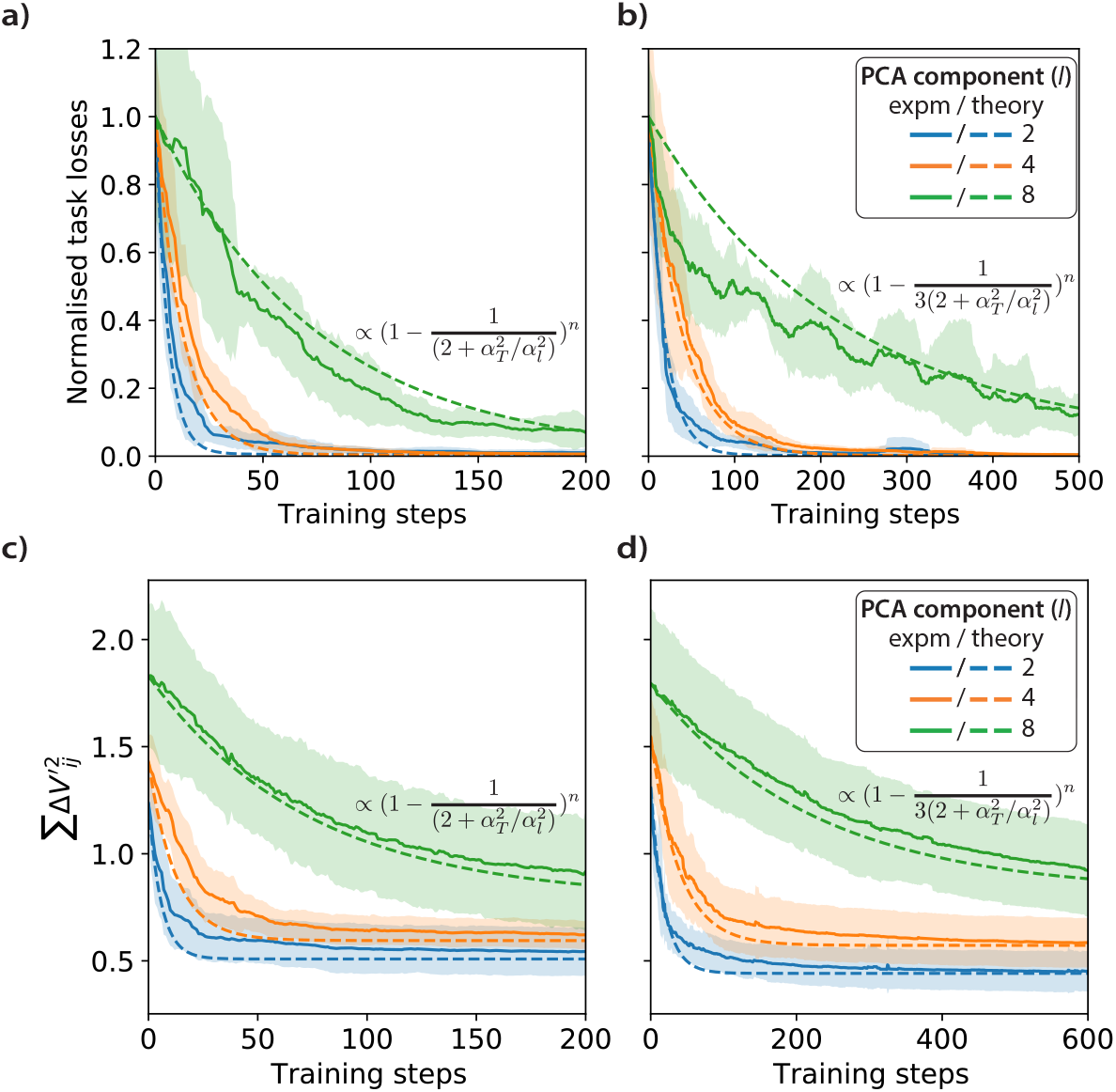
Example a) backpropagation and b) node perturbation dynamics for learning a single-dimensional cursor output based on a projection *U* onto different principal component numbers. For target principal components with small variances (increasing component number), the learning rate is significantly suppressed. Overlaid are predictions based on Eqns 21 and 22. c) & d) show the dynamics of 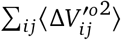 for the same experiments.

#### 4.1. Scaling of learning dynamics with number of output neurons

As noted in App. 2.1, for orthogonal BCI readout *U* projections, the learning dynamics of both backpropagation and node perturbation are insensitive to the number of output neurons (but not the target dimensionality). Since this is the case for both a very targeted credit assignment mechanism (backpropagation) and a high-variance but simple mechanism (node perturbation), this may be a common feature of many plausible neural credit assignment strategies. This suggests that a BCI task involving controlling a *single* neuron may have the same learning dynamics as learning to control a *population* of neurons, as long as both activities are projected to the same target dimensionality.

#### 4.2. Non-orthogonal BCI projections

In our derivation in App. 2.1 and in the rest of the paper, we assume that the BCI readout *U* projections are orthogonal. This is likely to be the case in most experiments. In the following we consider non-orthogonal *U* as an extension of the previous analysis, as this leads to quite different behaviour.

An interesting observation for non-orthonal *U* is that it can actually be beneficial to increase the number of neurons at the output layer *y* given a fixed target dimension *M*. For example, if *U* is randomly initialized, it will not be orthogonal or orthonormal (Fig. 10c). This leads to impaired learning due to the coupling between different elements of Δ*V* (Fig. 10a & b). So, in the case of random *U*, it is actually beneficial to increase the number of output neurons *M* given a fixed target *t* dimension.

**Figure 10.**
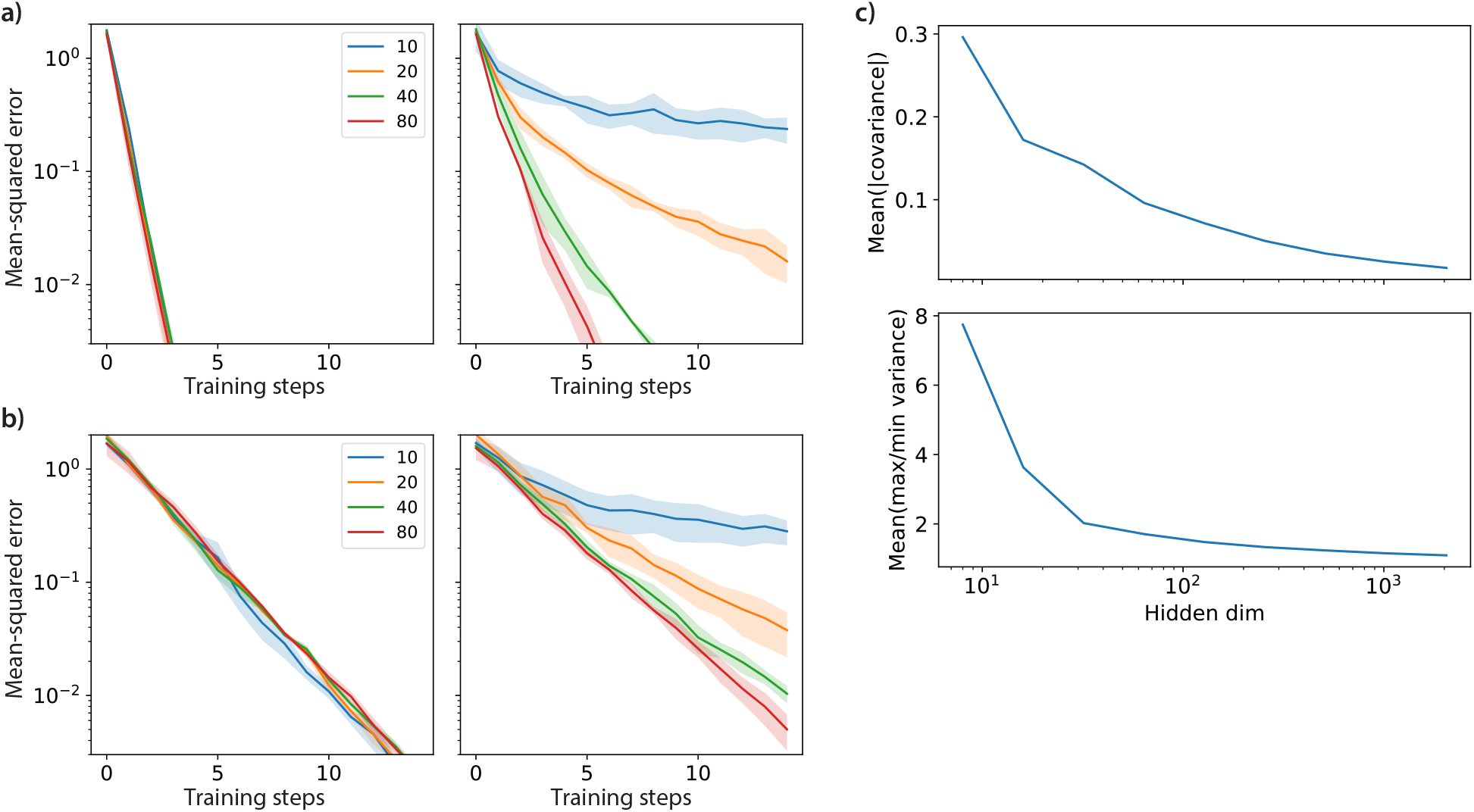
Learning dynamics under backpropagation (a) and node perturbation (b) for a BCI task with a random (left) or orthonormal (right) projection *U* as a function of the output layer *y* dimension. The dimension of the inputs *x* and BCI outputs *t* are both fixed at 10. The learning rate for orthogonal *U* is independent of the output dimension, while for a random matrix the learning rate actually increases with output dimension for both backpropagation and node perturbation. c) This increase is because random matrices have significant correlations. Here we generate random matrices of dimension (10, *N*) and measure the average covariances between the 10 output dimension projections, along with the mean ratio between the largest and smallest variances u^2^. Both will impact learning dynamics. As can be seen, larger output dimensionalities lead more orthonormal *U*.

The experimenter-controlled *U* in a BCI experiment is typically orthogonal, however, a non-orthogonal *U* could reflect a situation in which it is not the neurons at layer *y* that undergo learning, but a set of neurons in an upstream region, such that *U* is a composite transformation consisting of downstream connections as well as the BCI interface. In this case, recording from more BCI neurons could lead to an increase the number of neurons in this upstream learning region that have a causal influence on the BCI neurons, and hence could lead to a change in the rate of learning.

## Footnotes

1 In machine learning, gradient-based algorithms are typically assumed to *descend* gradients in order to minimise a loss. Alternatively, gradient ascent to maximise an objective can be assumed – such algorithms are often referred to as “hill-climbing”. In this study we will assume gradient descent.

2 Our theory results suggest that our conclusions are insensitive to the specific network layer used to simulate BCI experiments.

3 Note that, in contrast to our earlier experiments, these changes are not necessarily out-of-manifold. The initial distribution occupies a volume within the manifold – new activity patterns can also be within this manifold without being in-distribution.

